# Peptides at vesicle and mineral prebiotic interfaces

**DOI:** 10.1101/2025.06.13.659467

**Authors:** Ivan Cherepashuk, Mikhail Makarov, Radko Soucek, Vaclav Verner, Robin Krystufek, Romana Hadravova, Valerio Guido Giacobelli, Kosuke Fujishima, Sean F. Jordan, Liam M. Longo, Klara Hlouchova

**Affiliations:** Department of Cell Biology, Faculty of Science, Charles University, Prague 12800, Czech Republic; Institute of Organic Chemistry and Biochemistry, Czech Academy of Sciences, Prague 16610, Czech Republic; Earth-Life Science Institute, Institute of Science Tokyo, Tokyo 1528550, Japan; Graduate School of Media and Governance, Keio University, Fujisawa 2520882, Japan; Life Sciences Institute, School of Chemical Sciences, Dublin City University, Glasnevin, Dublin, 9, Ireland; Blue Marble Space Institute of Science, Seattle, Washington 98104, USA

**Keywords:** origins of life, peptides, minerals, vesicles, prebiotic compartments, amino acid alphabet evolution

## Abstract

The origin of life likely involved a complex interplay between organic molecules and mineral surfaces, yet the molecular details of these interactions remain poorly understood. Over recent decades, considerable research has focused on the individual roles of key biomolecules - such as RNA, lipids, and proteins - in early abiogenesis. However, this reductionist view offers only a partial picture because the emergence of life likely involved networks of molecular interactions that collectively shaped early functional assemblies. In this study, we examine the ability of peptides - arguably one of the most abundant early polymers - to interact with mineral surfaces and lipid vesicles, prebiotic interfaces and compartments. Using peptide libraries constructed from either prebiotically plausible or contemporary amino acids, *we demonstrate that while acidic residues drive peptide binding to mineral surfaces (such as fluorapatite, studied here), the inclusion of arginine - a basic residue that may have been accessible in specific prebiotic environments – synergistically enhances the mobilization of bioavailable phosphate from geological reservoirs*. Furthermore, we observe a functional divergence in vesicle interactions: while prebiotic alphabets promote dynamic membrane behaviours such as budding, libraries with ‘late’ canonical amino acids can help preserve vesicle integrity against salt-induced collapse. Our finding supports the view that interactions with peptides can elicit changes in both prebiotic minerals and vesicles, underscoring the importance of studying these systems collectively.

## Introduction

Mineral pores, coacervates, and vesicles (*enclosures bound by a lipid bilayer*) are believed to have been among the earliest compartmentalization structures on Earth. These structures are thought to have supported the emergence of life by influencing molecular assembly and stability, and serving as catalysts (Aquino et al., 2004; Biondi et al., 2007; Deamer et al., 2002; Lambert, 2008; Morigaki and Walde, 2007; Oparin, 1924; Rimola et al., 2013; West et al., 2017; Yu et al., 2013). While extensive research has explored the physicochemical properties of putative early life compartments, *including their formation, stability, and growth requirements* (Chen and Szostak, 2004; Cornell et al., 2019; Hentrich and Szostak, 2014), significantly less is known about their interactions with peptides. Yet, binding of peptides to mineral surfaces and vesicles may have a profound impact on their stability and catalytic properties. This knowledge gap reflects a long-standing divide in origin-of-life research, where compartmentalization, peptide chemistry, and mineral catalysis have often been investigated separately rather than as interconnected processes (Preiner et al., 2020).

Peptides were likely the most abundant and functionally versatile polymers in prebiotic environments (Fried et al., 2022). *Multiple studies have demonstrated the prebiotic accessibility of simple peptides, and ∼15-residue oligopeptides have been produced by a range of prebiotically relevant processes (Foden et al., 2020; Frenkel-Pinter et al., 2020; Kwiatkowski et al., 2021; Mayer et al., 2018).* Various lines of evidence demonstrate that even short peptides can play functionally significant roles: For example, di- and tri-peptides have been shown to catalyse the hydrolysis of proteins, nucleic acids, and esters (Cohen et al., 2024; Cornell et al., 2019; Gorlero et al., 2009). Amino acids and short peptides are also able to bind to simple fatty acid vesicles, resulting in an increase in vesicle stability to metal ions and high concentrations of salt (Cornell et al., 2019; Dalai and Sahai, 2019) and greater lamellarity (Cornell et al., 2019). Likewise, the presence of acidic residues can promote the binding of peptides onto mineral surfaces (Hoff et al., 2022). Taken together, these studies suggest that peptides were active participants in the origin of life, even before the emergence of templated protein synthesis.

Only about half of the canonical amino acids were likely to have been abundant on the early Earth or producible by a primitive metabolism. These “early canonical” amino acids, inferred from biosynthetic complexity, atmospheric discharge experiments, meteorite/asteroid analyses, and hydrothermal vent studies (Cleaves, 2010; Copley et al., 2005; Higgs and Pudritz, 2009; Trifonov, 2000), are listed in **Figure 1** as the 10E set. In addition to the 10E set, a plethora of non-canonical amino acids have been detected in prebiotic chemistry studies, and it is likely that both canonical and non-canonical amino acids were incorporated into primitive peptides (Zaia et al., 2008). Recent work has strongly favoured selection for protein foldability as the driving force for amino acid incorporation into the genetic code (Giacobelli et al., 2022; Hlouchová, 2024; Longo et al., 2020; Makarov et al., 2023; Solis, 2019; Tretyachenko et al., 2022). Here, we explore how primitive amino acid alphabets may have structured (or been structured by) early peptide-mineral and peptide-vesicle interactions. As the first peptides likely predated templated synthesis and had highly statistical sequences, we have chosen to employ random-sequence peptide libraries (**Figure 1**), which can better capture global differences between alphabets.

**Figure 1.**
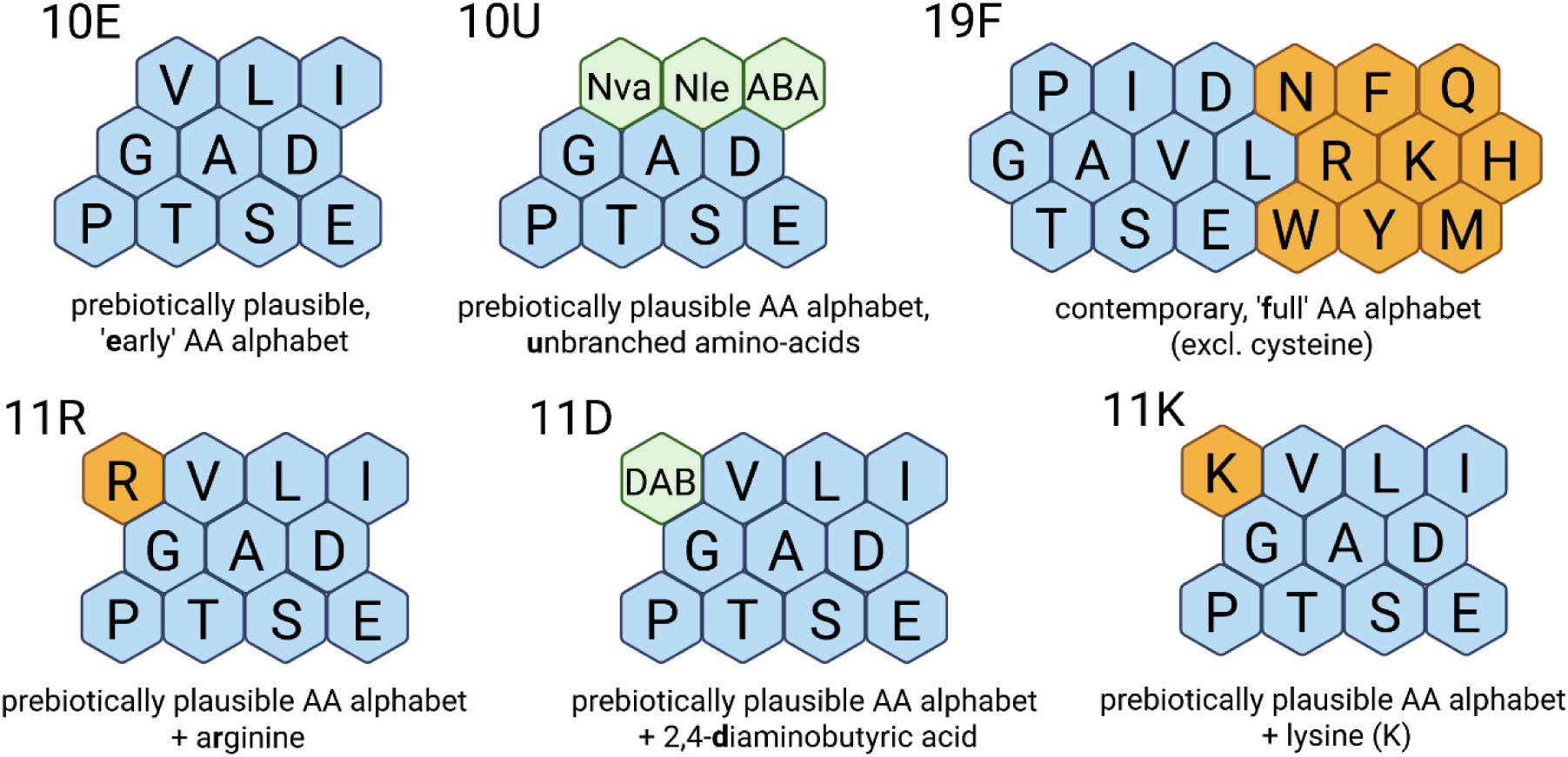
Amino acid composition of the random octapeptide libraries. Canonical amino acids are indicated using single letter codes. Blue hexagons denote prebiotically available amino acids (Higgs and Pudritz, 2009); orange hexagons denote “late” amino acids that require more elaborate biosynthesis; green hexagons denote prebiotically available amino acids that were not incorporated into the genetic code (non-canonical amino acids). DAB, Nva, NIe, ABA stand for 2,4-diaminobutyric acid, norvaline, norleucine, and α-aminobutyric acid, respectively.

The presence of phosphate minerals and clays at the origin of life is virtually uncontested. Decades of research into prebiotic membrane structures suggest that vesicles were also present on the early Earth (Deamer and Pashley, 1989; Gebicki and Hicks, 1976; Hanczyc et al., 2003; Hargreaves and Deamer, 1978; Jordan et al., 2019; Monnard and Deamer, 2011). *Membrane-forming molecules, such as single-chain amphiphiles, can be readily produced by Fischer-Tropsch chemistry (Cohen et al., 2023; McCollom et al., 1999), particularly in hydrothermal systems, and these molecules can be concentrated by association with mineral surfaces (Sahai et al., 2017). Moreover, the minimal fatty acid concentration required for vesicle formation can be as low as the micromolar range (Jordan et al., 2019). The observation that organic solvent-solubilized fractions of the Murchison meteorite self-assemble into vesicles in water (Deamer and Pashley, 1989) provides additional support for the view that different early Earth conditions could promote vesicle formation*.

Here, we synthesize six random octapeptide libraries to explore the potential role of peptide-mineral and peptide-vesicle interactions early in life’s history. We consider the interactions of peptides built from six different amino acid alphabets – five composed of prebiotically plausible amino acids and one reflecting a contemporary protein composition – with the ubiquitous phosphate mineral fluorapatite and simple decanoic acid:decanol (DA:DOH) vesicles. We find that primitive peptide libraries preferentially interact with the mineral surface and promote release of orthophosphate. These effects are likely due to an enrichment in acidic amino acids in the primitive amino acid alphabet. With respect to vesicles, we find that later amino acid alphabets exert slightly stronger vesicle-stabilizing effects in the presence of mono- and di-cations. We conclude that the bulk properties of primitive peptides were likely well-suited for mineral interaction.

## Materials and Methods

### Chemicals

For the solid-phase synthesis of combinatorial peptide libraries, Fmoc-Rink Amide AM resin (100-200 mesh, 0.64 mmol/g loading) and Fmoc-protected amino acids were purchased from Iris Biotech (Germany). N,N’-dimethylformamide (DMF) was purchased from Carl Roth (Germany), dichloromethane and diethyl ether were purchased from PENTA (Czech Republic). Ceramic hydroxyapatite and fluorapatite (type II, 40 μm particle size) were purchased from Bio-Rad (USA). All other chemicals were purchased from Sigma Aldrich (USA). For experiments related to vesicle interactions, Rhodamine 6G, DA were purchased from ThermoFisher Scientific (USA), and DOH was purchased from Sigma Aldrich.

### Chemical synthesis of combinatorial peptide libraries

8-mer combinatorial peptide libraries were prepared manually using Fmoc-based solid-phase synthesis. Synthesis of the peptide mixtures was accomplished by coupling Fmoc-protected amino acids individually to the equal aliquots of the solid support that were merged after coupling (split and mix approach). Prior to synthesis, Fmoc-amino acid/Oxyma Pure stock solutions were prepared by simultaneously dissolving Fmoc-amino acids and Oxyma Pure to the final concentrations of 300 mM and 375 mM in DMF, respectively. 500 mg of Rink Amide AM resin (0.32 mmol) was swelled in 20 ml of DMF for 1 h and subsequently deprotected by incubation with 15 ml of 20 % (v/v) piperidine in DMF for 1 h at room temperature. The resin was washed four times with 10 ml of DMF, thoroughly resuspended in 10 ml of DMF and split into equal aliquots, depending on the number of amino acids (10 for “10E”, 11 for “10U”, “11R”, “11D”, “11K”, and 19 for “19F” peptide libraries). For the synthesis of “10E” peptide library, 1.1 ml of Fmoc-amino acid/Oxyma Pure solutions and 64 µl of N,N′-diisopropylcarbodiimide were added to the resin aliquots, and the reaction mixture was incubated overnight at room temperature. The amounts of Fmoc-amino acid/Oxyma Pure solution and N,N′-diisopropylcarbodiimide used for coupling were scaled down for a higher number of amino acids (resin aliquots). Following the coupling, the resin aliquots were merged, washed five times with 10 ml of DMF and deprotected by incubation with 15 ml of 20% (v/v) piperidine in DMF for 1 hour at room temperature. The amino acid coupling and deprotection steps were performed eight times. All separation and washing steps were performed using 15 ml polypropylene syringes fitted with polypropylene frits. After the final coupling/deprotection step, the resin was washed five times with 10 ml of dichloromethane and dried *in vacuo*. The peptide mixture was cleaved off the dried resin by incubation with 10 ml of trifluoroacetic acid-triisopropylsilane-water (95:2.5:2.5, v/v) mixture for 2 hours at room temperature. The resin was filtered off, and the peptide mixture was precipitated with 150 ml of diethyl ether cooled down to 4 ℃ and submerged in liquid nitrogen for 10 min to maximize precipitation. The peptide mixture was subsequently centrifuged at 5000× g for 5 min at 4 ℃, and the peptide pellet was collected and lyophilized three times with 1 mM HCl overnight.

Additionally, 19F, 10E, and 11R libraries were synthesized with the inclusion of the TAMRA fluorescent dye for peptide co-localisation assays. Labelling was performed by overnight treatment with 3 eq. of 5(6)-carboxytetramethylrhodamine, 3.75 eq. Oxyma Pure and 3.75 eq. N,N′-diisopropylcarbodiimide in DMF after the final coupling/deprotection step.

### Quality control of combinatorial peptide libraries

The molecular weight distributions of 8-mer combinatorial peptide libraries were confirmed by mass spectrometry using UltrafleXtreme MALDI-TOF/TOF mass spectrometer (Bruker Daltonics, Germany) according to the standard procedure. Prior to the MALDI-TOF analysis, peptide libraries were dissolved in acetonitrile:water (50:50, v/v) mixture and mixed with 2,5-dihydroxybenzoic acid (DHB) that was used as a matrix.

The amino acid composition of 8-mer combinatorial peptide libraries was estimated by HPLC amino acid analysis performed on Biochrom 30 system (Biochrom, United Kingdom) using automated ninhydrin derivatization. Prior to the HPLC amino acid analysis, the library samples were hydrolyzed in 6 M HCl at 110 °C for 20 h with the hydrolysate evaporated and reconstituted with 0.1 M HCl.

### HPLC amino acid and mass spectrometry analyses of mineral binders from 8-mer combinatorial peptide libraries

10 mg of fluorapatite was gently rotated with 0.5 ml of 1 mg/ml peptide library solution in HMA buffer (20 mM HEPES, 20 mM MES, 20 mM sodium acetate, pH 5.5) for 21 hours at room temperature. The mixture was centrifuged at 15,000× g for 3 min at room temperature to separate apatite from the peptide library solution, and the supernatant was subsequently discarded. The apatite was resuspended in 0.5 ml of HMA buffer (pH 5.5), incubated for 3 min at room temperature and centrifuged at 15000× g for 3 min. The washing step was repeated four times. The bound peptides were subsequently released from apatite by overnight incubation with 220 μl of 50 mM HCl at room temperature. Following the incubation, the mixture was centrifuged at 15000× g for 3 min, and 150 μl of supernatant was submitted to MALDI-TOF or HPLC amino acid analyses. Mass spectrometry was performed once, an HPLC analysis was performed six times.

### Phosphate release by 8-mer combinatorial peptide libraries

Solubilization of fluorapatite by combinatorial peptide libraries at the pH of 5.5 and temperature of 55 ℃ conditions was estimated by molybdenum blue spectrophotometric assay (Murphy and Riley, 1962). Prior to analysis, molybdenum blue reagent was freshly prepared by mixing 2.5 M sulphuric acid, ammonium molybdate (40 g/l), 0.1 M ascorbic acid, and potassium antimonyl tartrate (1 mg Sb/ml) in 10:3:6:1 (v/v) ratio. 10 mg of apatite was gently rotated with 0.5 ml of 1 mg/ml peptide library solution in HMA buffer for 21 hours. The mixture was centrifuged at maximum speed (21.300× g) for 20 min to separate apatite from the peptide library solution, and 300 μl of supernatant was transferred to a new tube. The supernatant was centrifuged again at maximum speed (21.300× g) for 20 min to remove any residual apatite, and 250 μl of supernatant was transferred to a new tube. 5 μl of supernatant was mixed with MilliQ water to adjust the volume to 150 μl in a 96-well plate, and 30 μl of molybdenum blue reagent was added to each well. The reaction mixture was incubated for 20 min at room temperature, and the absorbance at 882 nm was measured using Infinite M Plex plate reader (TECAN, Switzerland). The amount of phosphate ions in the supernatant was quantified from the standard curve established using 2-fold serial dilutions of KH_2_PO_4_ solution in MilliQ water. The measurement was performed in technical triplicates. The analysis was performed three times.

### Preparation of stock decanoic acid:decanol (DA:DOH) vesicle solution

Decanoic acid (DA) and decanol (DOH) offer relative stability for vesicle experimentation and were observed in meteorite materials ^[6]^. All solutions were pre-warmed to 50 ℃ using a water bath. DA was dissolved in 340 volumes of acetate-phosphate-borate buffer (ABP, pH 7.4) or 5 mg/mL peptide library solution in ABP (in experiments where peptide libraries were present in the lipid solution prior vesicle formation). The mixture was mixed vigorously for 10 seconds, followed by the addition of 180 mM NaOH to achieve a DA concentration of 53 mM. DOH was then added in an equimolar concentration to DA, and the solution was mixed vigorously for 60 seconds to allow vesicle formation. *The pH was adjusted to 7.4 using HCl, resulting in a final concentration of 37.8 mM of DA and 38.4 mM of DOH*. The stock solution was then used in experiments downstream.

Preparation of peptide library solution for experimentation on DA:DOH vesicles. 5 mg of peptide library was dissolved in ABP to a final concentration of 5 mg/ml. This solution, freshly upon mixing by pipetting, was either added to DA (in experiments where peptide libraries were present in the lipid solution prior vesicle formation) or diluted with ABP to a concentration of 2 mg/ml. The stock solutions were used in consequent experiments shortly upon mixing by pipetting to ensure uniform peptide concentration.

Amino acid analysis of peptide libraries upon incubation with DA:DOH vesicles. 150 μl of the DA:DOH vesicle stock solution was mixed with 150 μl of 2 mg/ml solution of peptide libraries in ABP (“vesicle incubation” condition) or ABP alone (negative control) and incubated at 25 ℃ with shaking at 1000 RPM, 2 mm orbit for 1 hour. Additionally, peptide libraries were incubated in the absence of vesicle stock solution (replaced with ABP) in parallel (“vesicle control” condition in order to elucidate peptide library binding to the Amicon centrifugal filter itself). 10 μl of the resulting solution was dried out and submitted to amino acid analysis (“unfiltered” condition), and another 250 μl of the were loaded into Amicon Ultra-4 3K centrifuge filters and centrifuged at 3000× g for 10 minutes. 10 μl of the flow-through from the Amicon centrifuge filters were also dried out and submitted to amino acid analysis (“filtered” condition). Amino acid analysis was performed on a Biochrom 30 amino acid analyzer. Results from the negative control were subtracted from the results, and both relative concentrations of individual amino acids and whole peptide library concentration were compared relatively within each condition. All experiments were conducted in 6 replicates.

### Zeta-potential measurements

Measurements of zeta-potential of vesicle populations were conducted using Zetasizer Nano ZS90 (Malvern Panalytical) with DTS1070 cuvettes at standard manufacturer settings at 25 ℃. 350 μl of vesicle stock were mixed with 350 μl of either ABP or 2 mg/ml peptide library solution in ABP, and 70 μl of 4M NaCl or 100 mM MgCl_2_ solutions were also added, depending on the experiment, to a final concentration of 363 and 8.9 mM, respectively. Experiments were conducted in technical triplicates.

### Dynamic light scattering

Dynamic light scattering measurements were conducted using Zetasizer Nano ZS90 (Malvern Panalytical) with ZEN2112 cuvettes at 25 ℃. For this, 50 μl of vesicle-peptide solution was mixed with 50 μl of ABP, and 10 μl of 4M NaCl or 100 mM MgCl_2_ solutions were also added, depending on the experiment, to a final concentration of 363 and 8.9 mM, respectively. Side scattering measurements were then analysed to determine the size distribution of the samples. The experiment was conducted in technical triplicates.

### Spinning-disc confocal microscopy

50 μl of stock vesicle solutions were mixed in a 1:1 ratio with 2 mg/mL solutions of peptide libraries in ABP buffer on Glass Bottom 18-Well µ-Slides (Ibidi, Germany). Rhodamine 6G (excitation peak at 561 nm) was added to a final concentration of 1 µM. Depending on the experiment, either 10 μl of 4M NaCl solution was added to a final concentration of 363 mM or 10 μl of 100 mM MgCl_2_ solution to a final concentration of 9.1 mM. In alternative setups, 2 mg/ml of peptide library solution in ABP conjugated with TAMRA fluorescent dye (excitation at 561 nm) were mixed with their non-fluorescent counterparts (2 mg/ml in ABP) in a 1:50 ratio. In such instances, DiD fluorescent dye (excitation at 638 nm) was used for membrane imaging instead of Rhodamine 6G, which has an overlapping excitation range with TAMRA. Imaging was performed using a Nikon CSU-W1 spinning-disc confocal microscope with a 100× oil-immersion objective and a PRIME BSI camera (Teledyne Photometrics) at room temperature. Experiments were conducted three times.

### Transmission electron microscopy

100 μl of DA:DOH vesicle solution was diluted in 900 μl PBS buffer to a final concentration of 7.7 mM. Vesicle sample was then analysed in PBS buffer by negative staining. Parlodion-carbon-coated grids were floated on the top of 10 μl drop of the sample for 5 min. Then the grids were stained of 2% uranyl acetate, for 2 × 30 s and dried. The grids were examined with a JEOL JEM-2100 Plus electron microscope operated at 200 kV (JEOL Ltd., Japan).

### Turbidity measurement

Turbidity measurements were conducted in 96-well plates on a microplate reader (Infinite M Plex, Tecan, Switzerland). Vesicle solutions were prepared in presence of peptide libraries with free fatty acids as outlined above. 10 μl of 4 M NaCl or 10 μl of 100 mM MgCl_2_ solutions were added to final concentrations of 363 mM or 9.1 mM, respectively. OD_490_ measurement was then conducted in triplicates, and absorbance was compared against average vesicle sample absorbance at the absence of peptide libraries, which was put at 100% for every salt addition condition screened (control (no salts added), 363 mM NaCl or 9.1 mM MgCl_2_).

### Critical vesicle concentration measurement

Critical vesicle concentration (CVC) was established by preparing serial dilution of vesicle stock solution with or without peptide libraries (from 0.35 to 11.33 mM, 100 μl final sample volume in a 96-well plate), adding 2 μL of pyrene dye (pure, 1 mM stock solution in methanol) to final concentration of 5 μM and measuring fluorescence emission at 384 and 373 nm using Infinite M Plex microplate reader (Tecan, Switzerland) at 25 ℃. The 384/373 ratio curve was then plotted against vesicle concentration and the approximation of CVC was evaluated at the 1.15 384/373 ratio cutoff point. The experiment was conducted in triplicates.

### Statistical analysis

All data were analyzed using Python (version 3.11) with numpy, pandas, seaborn, scipy, and matplotlib libraries. Normality of distributions was assessed using the Shapiro–Wilk test. Pairwise comparisons between the Control group and peptide libraries were performed using the non-parametric Mann–Whitney U test. To account for multiple testing, Holm-Bonferroni correction was applied to adjust the p-values. Differences were considered statistically significant at p < 0.05 after correction. For CVC analysis, *group differences were assessed using a Kruskal–Wallis test followed by Holm-corrected Mann–Whitney post hoc comparisons. No statistically significant differences were detected (p = 0.10)*.

## Results and Discussion

### Peptide libraries: properties and motivations

Six different octapeptide libraries with fully random sequences were synthesized (**Figure 1**). Amino acids were selected based on their prebiotic plausibility (Higgs and Pudritz, 2009) and biophysical properties (Makarov et al., 2023): The 10E library consists of the ten early canonical amino acids (Longo et al., 2013). The 10U library replaces the three canonical branched amino acids (Val, Lue and Ile) with three prebiotically available, unbranched amino acids (norvaline, norleucine, and alpha-aminobutyric acid). Thus, by comparing the 10E and 10U libraries, the effect of amino acid branching, and the reduction in side-chain entropy it entails, can be assessed.

Both the 10E and 10U libraries are devoid of positively charged amino acids, making the N-terminus the only positively charged group on these peptides (McDonald and Storrie-Lombardi, 2010). *To study the role of basic amino acids, three alphabets based on the early canonicals (10E), with the addition of a single basic amino acid type, were constructed: 11D (addition of 2,4-diaminobutyric acid or DAB), 11R (addition of Arg), and 11K (additional of Lys).* DAB was likely present in prebiotic environments (Zaia et al., 2008) and is hypothesized to have been the first coded basic amino acid, present during an intermediate stage of the genetic code (Copley et al., 2005). Arg, although not typically detected in meteorites or atmospheric discharge experiments, may have been accessible via a cyanosulfidic protometabolism (Patel et al., 2015). In addition, conversion of prebiotically available ornithine into Arg by guanidination has been demonstrated, both for the free amino acid and for ornithine-containing prebiotic peptides (Ariely et al., 1966; Longo et al., 2020). Lys is hypothesized to be the most recent basic amino acid addition to the genetic code (Trifonov, 2000).

Finally, as a point of comparison to the primarily prebiotic alphabets above, the 19F library – which includes all canonical amino acids except Cys – was constructed. The exclusion of Cys was due to limitations of Fmoc-based peptide synthesis. Unlike the other libraries, 19F includes aromatic amino acids (Phe, Tyr and Trp) and amide-containing side chains (Asn, Gln). Note also the presence of His, which is often involved in metal binding (Li et al., 2024).

In each case, the amino acid types are approximately uniformly distributed within a peptide library (**Figure S1**) and not biased by either prebiotic availability or frequency in proteins. Given the large gaps in our understanding of primitive peptide synthesis, this decision is meant to retain as much neutrality as possible. Assuming a perfectly uniform distribution of amino acids, the properties of the bulk library can be estimated (Makarov et al., 2023): *the order of hydrophobicity, from highest to lowest, is 19F > 11K > 11R > 11D > 10U > 10E. The fraction of charged residues, from highest to lowest, is 11R (0.27) = 11K (0.27) = 11D (0.27) > 19F (0.26) > 10E (0.2) = 10U (0.2). Finally, the estimated median pI of each library is 10E (5.46) < 10U (5.49) < 11K (5.84) < 11D (5.85) < 11R (5.94) < 19F (6.13)*.

### Peptide-vesicle and peptide-mineral interactions are influenced by library composition

First, we measured the extent to which each of the octapeptide libraries interacts with mineral and vesicle surfaces. The mineral fluorapatite was selected due to its abundance and prebiotic availability, as well as its potential role as a prebiotic source of phosphate (Paytan and McLaughlin, 2007). Fluorapatite is more stable than hydroxyapatite and found on both Earth and Mars (Hausrath and Tschauner, 2013; Martínez et al., 2023). The n-decanoic acid and n-decanol (DA:DOH) vesicle system was selected due its putative prebiotic availability (Cohen et al., 2024; Cornell et al., 2019). *Peptide binding (**Figure 2A**) was measured after co-incubation with mineral powder or vesicles. Filtration (vesicles) and centrifugation (mineral grains) were used to separate the prebiotic substrate-bound peptides from the bulk solution. Finally, peptides samples were subjected to amino acid analysis (see **Methodology**)*.

**Figure 2.**
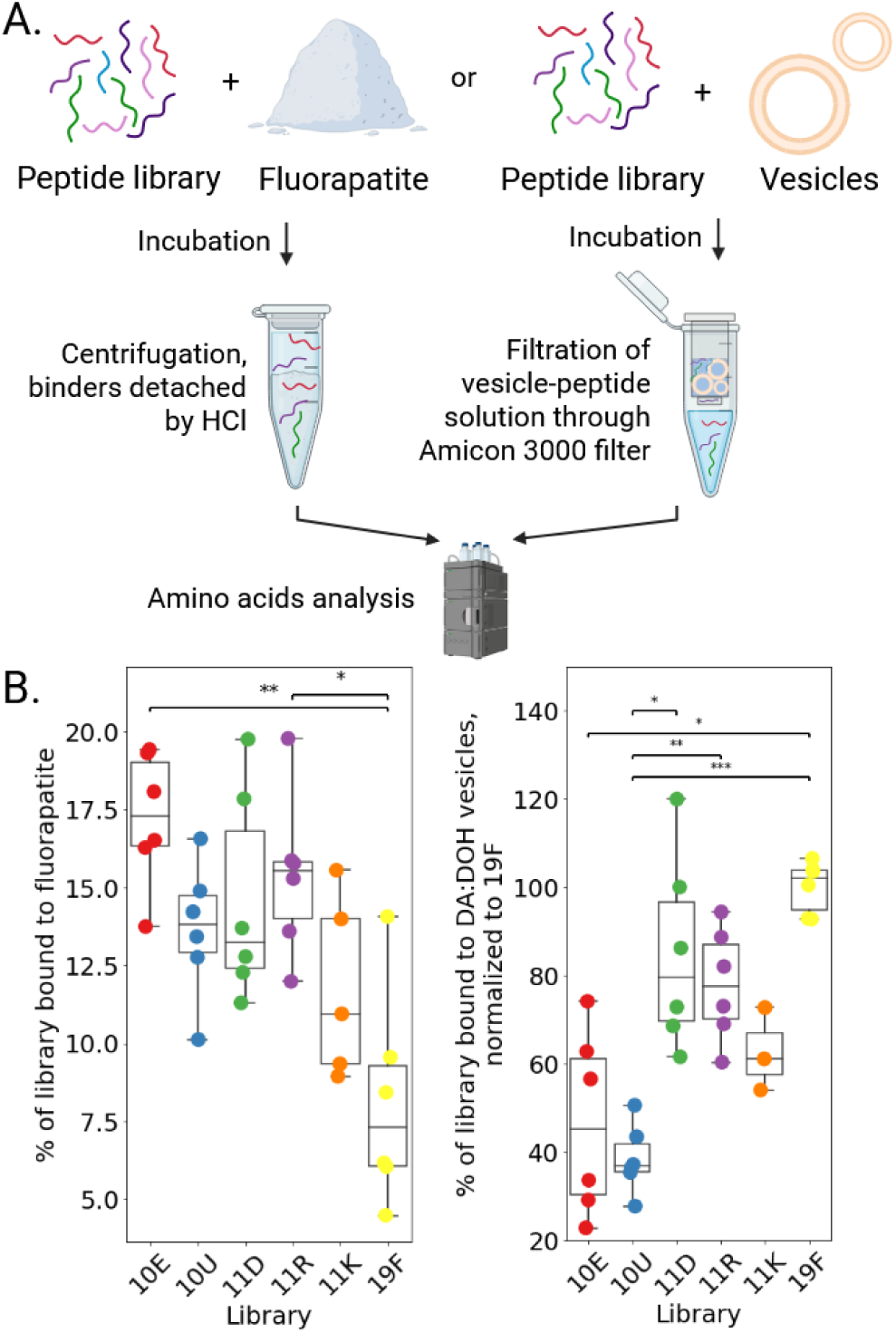
Peptide library association with prebiotic surfaces. A. Outline of the experimental pipeline. Since it is not feasible to centrifuge fatty acids down and alcohols interfere with absorbance at 220 nm, HPLC was used to establish the reduction in peptide concentration upon binding for both setups. B. Library binding to fluorapatite (left) or DA:DOH vesicles (right) calculated from amino acid analysis. Briefly, the amount of peptide bound to fluorapatite was estimated to be the sum of individual amino acid molar amounts in the sample. The same approach was taken in the vesicle-peptide assay but, due to limitations of the filtration-based assay, only relative concentrations are reported. Whiskers indicate the minimum and maximum values, boxes span the second to the third quartile, and the median value is indicated with a line. Statistical significance calculated using the Holm-Bonferroni-corrected U-test is indicated by asterisks: p < 0.05 (*), p < 0.01 (**), and p < 0.001 (***).

*Every peptide library screened exhibited some fluorapatite binding, with 5-20% of peptide molecules retained on the surface, as measured by an experiment in which bound peptides were desorbed from the surface of the mineral (****Figure 2A*** *and **2B**, left). We find that the peptide libraries based on 10E (10E, 10U, 11D, 11K, 11R) bind more strongly to fluorapatite than the 19F library (**Figure 2B**, left). In addition, amino acid analysis reveals that Asp and Glu residues were enriched in fluorapatite-bound peptides relative to the full library in each case (**Figure S2**).* These results are consistent with the known affinity of acidic residues for apatite, as acidic side chains can coordinate calcium ions (Hoff et al., 2022; Tavafoghi and Cerruti, 2016).

We note that several factors may influence the binding affinity of the peptide libraries to fluorapatite. Foremost is pH: Our experiments were performed at pH 5.5, where the surface of fluorapatite is expected to be positively charged based on a point of zero charge (PZC) of 6.8 (Bell et al., 1973). In practice, however, the fluorapatite interfacial charge tends to be more negative than that predicted by PZC (Chaïrat et al., 2007). By changing the pH, it may be possible to tune binding; however, as long as Asp and Glu can coordinate calcium, we expect acidic amino acids to be the primary mediators of binding. Second*, we note that higher sintering temperatures yield apatite samples with weaker peptide binding propensity (Jeyapalina et al., 2022). The fluorapatite used in our experiments was sintered at a temperature of 700 °C (Trzaskowska et al., 2023)*.

*Next, we evaluated the binding of the peptide libraries to DA:DOH vesicles (**Figure 2B**, right). We first confirmed the presence of vesicles using transmission electron microscopy and fluorescent imaging (**Figure S3**). After incubation, Amicon® Ultra-4 3K centrifugal filters, which use a regenerated cellulose membrane, were used to separate the vesicle-bound and unbound peptides (****Figure 2A****), as has been done previously (Black et al., 2013; Cornell et al., 2019). Although we observed considerable non-specific peptide binding to the filter membrane itself, comparisons with vesicle-free controls demonstrated that this effect was uniform across all libraries (**Figure S4**).* Because this background loss represents a systematic bias rather than a variable one, we report our findings as relative comparisons of vesicle affinity.

*We find that the 10E and 10U libraries have an 80% higher peptide concentration in the unbound phase relative to 19F than libraries with positively charged amino acids (11D, 11R, and 11K). This result suggests that positively charged amino acids play a role in peptide-vesicle association, in agreement with previous reports on basic amino acid behaviour (Cohen et al., 2024; Kamat et al., 2015; Kwiatkowski et al., 2021; Mitchell et al., 2000). Interestingly, the 11D library (containing DAB) displayed a similar degree of vesicle binding to the 11R (containing Arg) and 11K (containing Lys) libraries.* This result suggests that fatty acid vesicle binding may not have exerted a strong selective force on the amino acid alphabet, consistent with our earlier interpretation that foldability was the primary property under selection. Moreover, we do not find substantial differences in the amino acid composition of peptides filtered from the vesicle solution relative to the total library (**Figure S4**).

### Peptide libraries promote phosphate release from fluorapatite

As phosphate is essential for energy conservation, information storage, and encapsulation, we determined whether the peptide libraries could promote the release of orthophosphate from fluorapatite. To detect released orthophosphate, we employed the molybdenum blue assay (**Figure 3B**; see **Figure S5** for a representative standard curve). We find that all screened libraries induced orthophosphate release from fluorapatite, increasing the concentration of soluble orthophosphate by up to a factor of two relative to the mineral alone (**Figure 3C**). *Notably, this effect appears to be composition-dependent. While all libraries promoted dissolution, those containing Arg (the **11R** and **19F** libraries) consistently elicited the highest levels of phosphate release. We propose a cooperative mechanism driving this efficiency. All libraries contain Asp and Glu, which likely coordinate calcium ions, destabilizing the surface lattice via calcium leaching, thereby shifting the equilibrium toward dissolution (Matreux et al., 2025). However, the inclusion of Arg appears to synergize with this effect. The guanidinium group of Arg is unique in its ability to form stable, bidentate hydrogen bonds with phosphate and phosphonate groups (Woods and Ferré, 2005). This highlights a potential advantage of the ‘modern’ basic amino acid Arg over simpler prebiotic basic amino acids in accessing geochemical phosphate reservoirs*.

**Figure 3.**
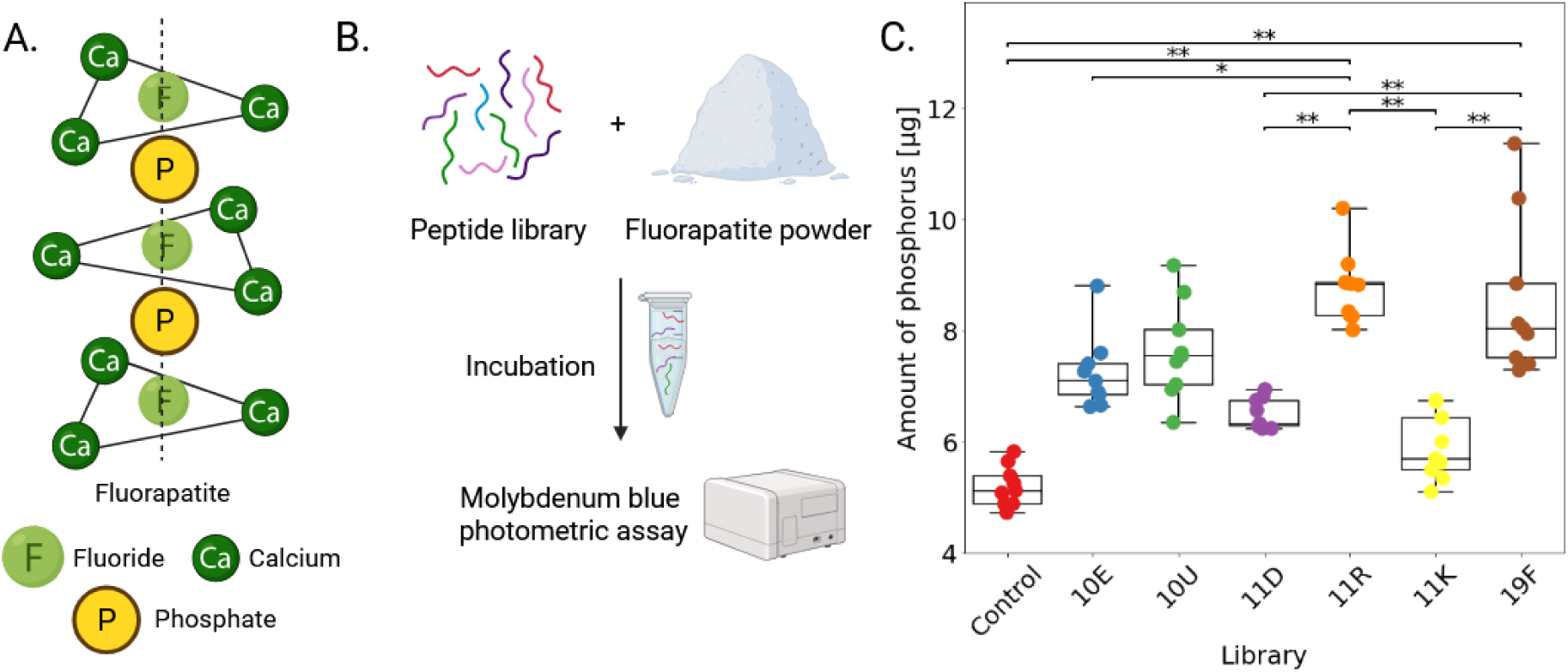
A. Chemical structure of fluorapatite Ca_10_(PO_4_)_6_F_2_ (Hoff et al., 2022; Posner, 1985). B. Experimental workflow for detecting orthophosphate release from fluorapatite. C. Orthophosphate release from fluorapatite. The “Control” sample lacks any peptides libraries, and incubation was performed in buffer (see **Methodology**). All peptide libraries except for 11K promoted significant orthophosphate release relative to the buffer control (p < 0.05 for all 5 comparisons to the control, as assessed by the Holm-Bonferroni-corrected U-test). Box and whisker plot as in Figure 2.

### Peptide libraries affect vesicle stability in high-salt conditions

We next determined whether there is a composition-dependent impact of peptide libraries on vesicle structure, stability, or aggregation. We address these questions using optical density at 490 nm (OD_490_; a measure of aggregation and lamellarity of fatty acid vesicles (Black et al., 2013; Cornell et al., 2019; Wang et al., 2019)), spinning-disc fluorescent microscopy, and dynamic light scattering (DLS) of vesicles created in the presence of peptide library solution. We find that vesicle morphology, OD_490_, and average particle size are largely unaffected by any of the six peptide libraries under no-salt conditions (**Table S1**). However, upon addition of 363 mM NaCl or 8.9 mM MgCl_2_, significant differences were observed.

We find a large increase in OD_490_ upon addition of MgCl_2_, if peptide libraries were present in the sample (**Figure 4A**). *This effect can be partially explained by the formation of punctate structures (Cornell et al., 2019) and/or multilamellar vesicles, depending on the peptide library used (**Figure 4B**, images O-U). We note, however, that the formation of Mg(OH)_2_ can also result in an increase in optical density. The destabilization of vesicles in the presence of both peptide libraries and divalent cations may originate from increased osmolarity or disruption of hydrogen bonding networks*. These effects seem to be negated in part by the late canonical amino acids present in the 19F peptide library.

**Figure 4.**
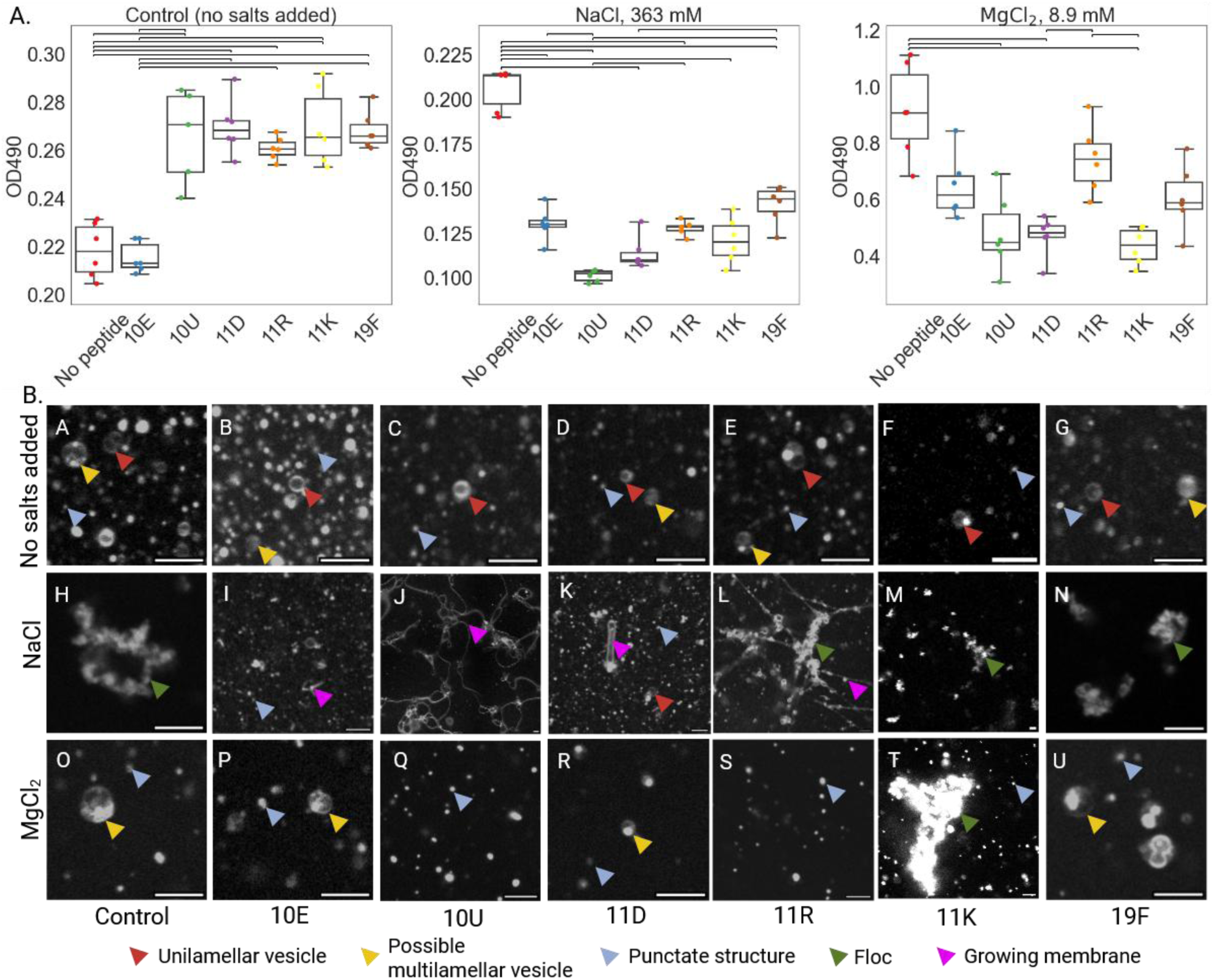
A. Optical density at 490 nm for vesicles-peptide library samples (“10E”, “10U”, “11D”, “11R”, “11K” and “19F”). Note that peptide libraries were added to free fatty acids before vesicle formation. Three conditions were tested: no salts added, NaCl (363 mM final) and MgCl_2_ (8.9 mM final). Box and whisker plot as in Figure 2. B. Micrographs of DA:DOH (1:1) vesicles under different conditions. Vesicular flocculation can be observed upon addition of NaCl. A reduction in the number of visible particles occurs upon addition of MgCl_2_. Staining with Rhodamine 6G. Scale bar: 5 µm; Red arrow - unilamellar vesicle; yellow arrow - possible multilamellar vesicle; blue arrow - punctate structure; green arrow – flocculation, pink arrow - growing membranes.

Upon addition of NaCl to peptide-vesicle samples, we observe formation of filamentous structures that resemble growing and budding fatty acid membranes (Hentrich and Szostak, 2014) (**Figure S6**) in all but the 19F library. These filamentous structures are especially pronounced in the presence of the 11R, 11D, and 10U libraries (**Figure 4B**, images J-K). Curiously, these libraries also show the greatest disruption in vesicle formation upon addition of MgCl_2_ (**Figure 4B**, images P, Q, and S). The *19F library, despite not showing definitive protective effects against flocculation, seems to preserve vesicle integrity in the presence of MgCl_2_, as indicated by microscopically observable membranes (**Figure 4B**, images O and U) and an increase in OD_490_ compared to control conditions (which may report on the degree of lamellarity or the formation of Mg(OH)_2_; **Figure 4A**)*.

As surface charge influences vesicle-peptide interactions (Chen and Szostak, 2004), we calculated the ζ-potential of DA:OH vesicles in the presence and absence of salt. We find that 38.5 mM DA:DOH vesicles have a ζ-potential of −61.5±4.2 mV, which drops to −17.6±9.2 mV and −31±0.4 mV in the presence of 363 mM NaCl or 8.9 mM MgCl_2_, respectively (**Table S1**). This reduction in ζ-potential likely contributes to vesicle flocculation because vesicle surface charge is screened by the presence of ions, decreasing vesicle-vesicle repulsion. Addition of peptide libraries only slightly reduces the ζ-potential, with the greatest reduction observed for the 10U library, yielding a ζ-potential of - 49.9±0.9 mV. Thus, a change in ζ-potential upon peptide binding does not seem to mediate the effect of the peptide libraries on vesicle structure and stability.

*To test whether the peptide libraries alter membrane formation propensity, we measured the critical vesicle concentration (CVC) using pyrene fluorescence. Peptide libraries caused only a minor, non-significant (p = 0.10) increase in CVC from ∼1 mM to ∼1.2-1.4 mM of DA:DOH (calculated as a sum of molar concentrations of DA and DOH) or had no measurable impact on CVC (**Table S3**).* Thus, changes in vesicle stability cannot be explained by an increase in CVC.

Taken together, our results suggest that the 19F library is the most beneficial for vesicle stability in the presence of salts. This library is the most hydrophobic and least soluble (Makarov et al., 2023) of the tested libraries, while also having the most basic isoelectric point. The extent to which the benefits of 19F are due to general properties (hydrophobicity, charge) or specific interaction modes with one or several of the late amino acids is unclear. If vesicle stabilization is specific to the properties of the late amino acids, however, it may suggest a potential benefit to expanding the genetic code, though whether this would be to stabilize membranes or augment the interaction of peptides with membranes is similarly unclear. In addition, we find consistent differences between the 10E and 10U libraries, with 10U having indications of stronger vesicle binding. This difference may be due to the longer length of the unbranched sidechains of norvaline and norleucine relative to their branched counterparts, facilitating fatty acid packing and/or membrane penetration. *We hypothesize that peptides with positively charged and unbranched amino acids may have promoted lipid packing or cross-linking of lipid surfaces within primitive mixtures*.

### Addition of MgCl_2_ or NaCl enhances peptide library co-localisation with DA:DOH vesicles

Surface charge is of importance to the vesicle-peptide interactions and can mediate peptide binding to membrane surfaces (Chen and Szostak, 2004). We used ζ-potential measurements to assess the correlation between the vesicle behaviour and their relative charge. We find that 38.5 mM DA:DOH vesicles have a ζ-potential of −61.5±4.2 mV. In the presence of 363 mM NaCl or 8.9 mM MgCl_2_, the ζ-potential drops to −17.6±9.2 mV and −31±0.4 mV, respectively (**Table S1**). This reduction in ζ - potential likely contributes to vesicle flocculation because vesicle surface charge is negated by the presence of ions, decreasing vesicle-vesicle repulsion. Addition of peptide libraries only slightly reduces the ζ-potential,with the greatest reduction observed for the 10U library, yeilding a value of −49.9±0.9 mV. We argue that the ζ-potential might not be a decisive factor mediating the effect of the peptide libraries on vesicles.

To visualize peptide-vesicle co-localisation, we labelled peptide libraries with only canonical amino acids (19F, 11R, and 10E) with *5-carboxytetramethylrhodamine* (TAMRA) and imaged them using spinning-disc microscopy alongside *1,1’-dioctadecyl-3,3,3’,3’-tetramethylindodicarbocyanine* (DiD)-stained vesicles. We observed that even peptide libraries with weaker vesicle association in the filtration assays exhibited marked co-localization with vesicle membranes upon the addition of NaCl and Mg^2+^ (**Figure 5**), possibly due to the reduction in the ζ-potential of the vesicles. We propose that the neutralisation of repelling forces on both membranes and peptides by NaCl and Mg^2+^ increases co-localisation of peptides and fatty acids, and that Mg^2+^ can also act as a “bridging” agent between these molecules. We argue that this co-localisation of peptide libraries indicates potential association, binding, or even membrane integration, as recently reported by molecular dynamics simulation (Cohen et al., 2024).

**Figure 5.**
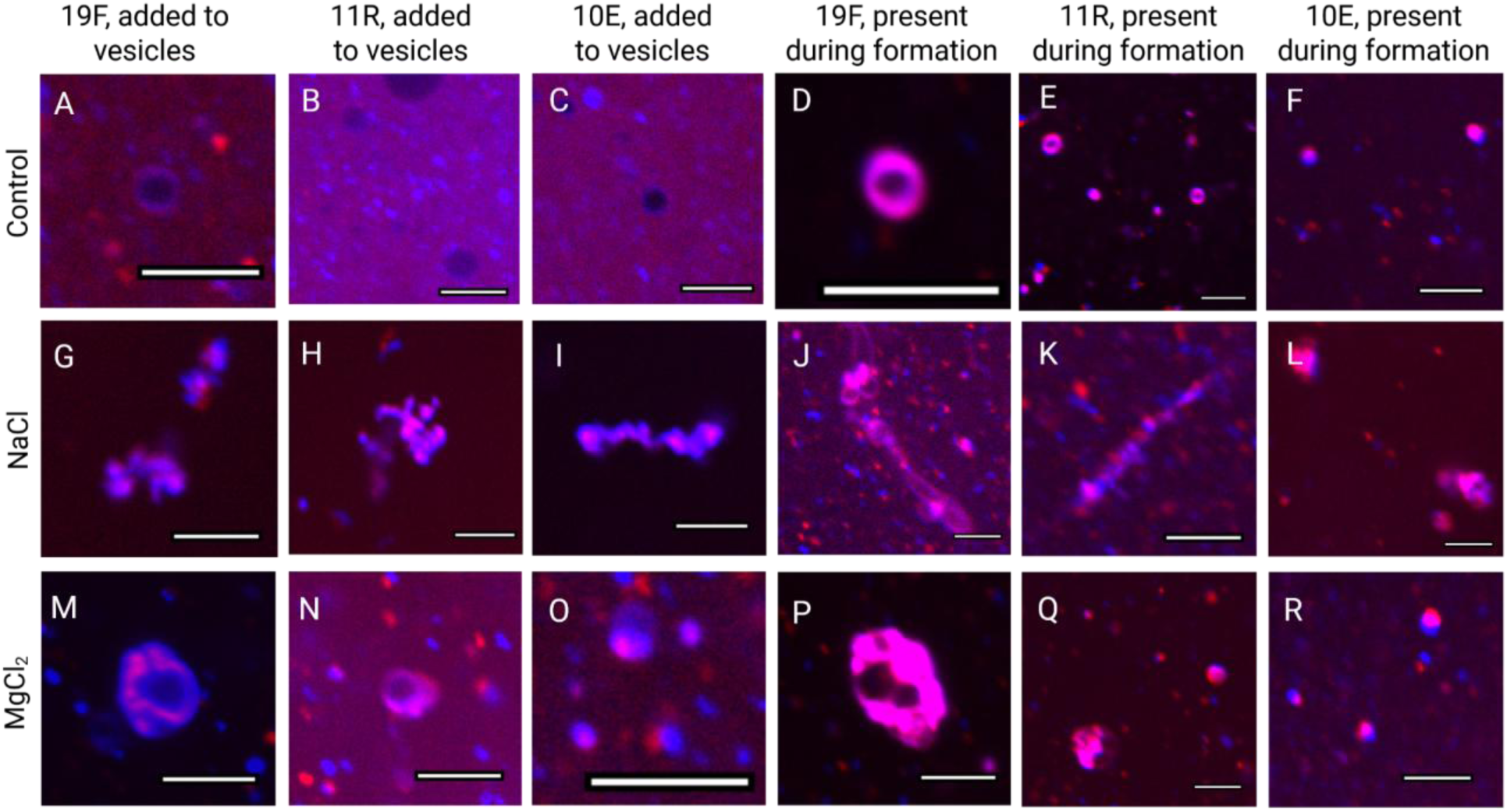
Fluorescence microscopy images of vesicles with peptide libraries either after (columns A-C, G-I, M-O) or before (columns D-F, J-L, P-R) vesicle formationunder three conditions: Control (top row), 363 mM NaCl (middle row), and 9.1 mM MgCl₂ (bottom row). Peptide libraries (19F, 11R, and 10E) are stained with TAMRA (red channel), while lipids are stained with DiD (blue channel). Addition of NaCl (middle row) induces vesicle aggregation and peptide co-localisation with the membranes (columns G-I), with similar, though less pronounced effects upon addition of MgCl_2_ (columns M-O). Peptide libraries present in the solution before vesicle formation (columns D-F, J-L, P-R) consistently exhibit more stable co-localisation with vesicular membranes across conditions. Scale bars represent 5 µm.

Notably, we find visual co-localisation of peptide libraries with DA:DOH membranes (but not within the vesicle lumen) only if peptide libraries were added to free fatty acids prior vesicle formation. When added to pre-formed vesicles, no observable co-localisation occurred between DA:DOH vesicles and 10E and 10R libraries. We suggest that peptide libraries integrated into the forming membranes and potentially modulate membrane formation, as indicated by the relative changes in OD_490_ in earlier experiments. In contrast, peptides added to vesicle post-assembly only weakly associated with the vesicle surface.

DA:DOH vesicles were reported to allow the passage of large organic molecules such as ribose and 5′-imidazole-activated dAMP through the membrane, measureable *on the timescale of hours* (Budin and Szostak, 2011). Despite this result, we did not find substantial accumulation of fluorescently labelled peptides in the vesicle lumen if peptides were added to pre-formed vesicles (**Figure 5**). We hypothesise that, due to inclusion of basic amino acids, 19F and 11R libraries could potentially permeate through the fatty acid membranes, but, due to the short time period in which the measurements were conducted upon the formation of vesicles (∼5-10 minutes), the membrane permeability should not have been substantial enough to observe.

### Limitations of the study

Our study is subject to several limitations. First, the libraries used here are a uniform mix of random peptides. As such, individual sequences with exceptional properties are not accessible to the experiments conducted in this work. This feature explains why the effects of the studied libraries are often consistent with the properties of individual constituent amino acids. Moreover, because our libraries contain a mixture of peptides, positive synergistic effects (that enhance the properties of the bulk library) as well as quenching effects (that reduce the properties of the bulk library) are possible. In the latter case, for example, interaction between positively and negatively charged peptides may compete with interaction of positive charged peptides with vesicles.

Both mineral surfaces and fatty acid vesicles are complex systems, and caution must be taken when interpreting experimental results. Fluorapatite binding and orthophosphate relase, for example, are likely influenced by the surface heterogeneity of the sample and the amount of impurity within the mineral. Geologically relevant sample flow, as achieved in flow chambers or similar setups, likely also impacts the properties of mineral-peptide systems (Matreux et al., 2025). Vesicles, on the other hand, are extremely sensitive to sample handling and environmental conditions. We find that minor changes in the pH of the final solution or the duration of sample shaking can lead to observable changes in vesicle stability (**Figure S7**). As such, quantitative descriptions of vesicle stability and morphology should be approached with caution. To mitigate these effects, the same vesicle stock solution was used for every condition screened here.

Lastly, our study exclusively explored only a single variant of prebiotically plausible vesicles and minerals as models for potential prebiotic interfaces. A great variety of possibilities exist for prebiotic peptide-interface interactions depending on the model mineral or membrane forming molecule(s) used.

### Conclusions

As plausible predecessors of proteins, peptides likely bridged the gap between prebiotic geochemistry and the emergence of biochemistry. In this study, we systematically explored how peptide composition influences interactions with two critical prebiotic interfaces: mineral surfaces and lipid vesicles.

We find that the ‘early’, prebiotically available amino acid repertoire was sufficient to mediate interactions with *fluorapatite* and promote phosphate release. While early peptide compositions promoted membrane dynamics (such as budding) in NaCl, they rendered vesicles susceptible to salt-induced instability. Libraries containing later amino acids, however, seemed to promote vesicle stability in the presence of salts.

## Acknowledgements

This work was supported by a Research Grant from the Human Frontiers Science Program (HFSP) to KH and KF (https://doi.org/10.52044/HFSP.RGEC272023.pc.gr.168579). MM and IC acknowledge support from Grant Schemes at Charles University (CZ.02.2.69/0.0/0.0/19_073/0016935, project no. START/SCI/148). KH, MM, and IC acknowledge the Biophysical techniques Core Facility of the Czech Infrastructure for Integrative Structural Biology Instruct-CZ Centre, supported by Ministry of Education, Youth and Sports of Czech Republic (MEYS CR) Infrastructure project LM2018127 and European Regional Development Fund-Project, “UP CIISB” (No. CZ.02.1.01/0.0/0.0/18_046/ 0015974). LML acknowledges support from HFSP (RGEC29/2025) and the NASA (80NSSC25K7873). SFJ acknowledges support from the European Research Council (ERC) under the European Union’s Horizon Europe research and innovation programme (grant agreement No 1101114969) and the Science Foundation Ireland (now Research Ireland) (SFI Pathway award 22/PATH-S/10692).

## Authorship confirmation/contribution statement

**Ivan Cherepashuk**: investigation, validation, formal analysis, writing - original draft, review & editing, visualisation (lead); conceptualisation, methodology, funding acquisition (equal).

**Mikhail Makarov**: conceptualisation, investigation, methodology, funding acquisition (equal).

**Radko Soucek**: investigation, resources, methodology (equal); writing - original draft, review & editing (equal).

**Václav Verner:** investigation, resources, methodology (supporting); writing - original draft, review & editing (supporting).

**Robin Krystufek**: resources, investigation (supporting).

**Romana Hadravova**: resources, investigation (supporting).

**Valerio Guido Giacobelli**: supervision, conceptualisation, project administration, writing - original draft, review & editing, visualisation (supporting).

**Kosuke Fujishima**: supervision, conceptualisation, project administration, funding acquisition, writing - review & editing (equal).

**Sean F. Jordan**: supervision, conceptualisation, methodology, writing - review & editing (equal).

**Liam M. Longo**: supervision, conceptualisation, methodology, writing - review & editing (lead).

**Klara Hlouchova**: supervision, conceptualisation, project administration, funding acquisition, writing - review & editing (lead).

## Authors’ disclosure statement

The authors declare that there are no conflicts of interest related to this work.

## Notes

### Competing Interest Statement

The authors have declared no competing interest.

### Summary of Updates

The manuscript has undergone major revisions upon submission to a journal. Quality of writing, data and figures was greatly improved.

