## Supplemental files for "Peptides at vesicle and mineral prebiotic interfaces"

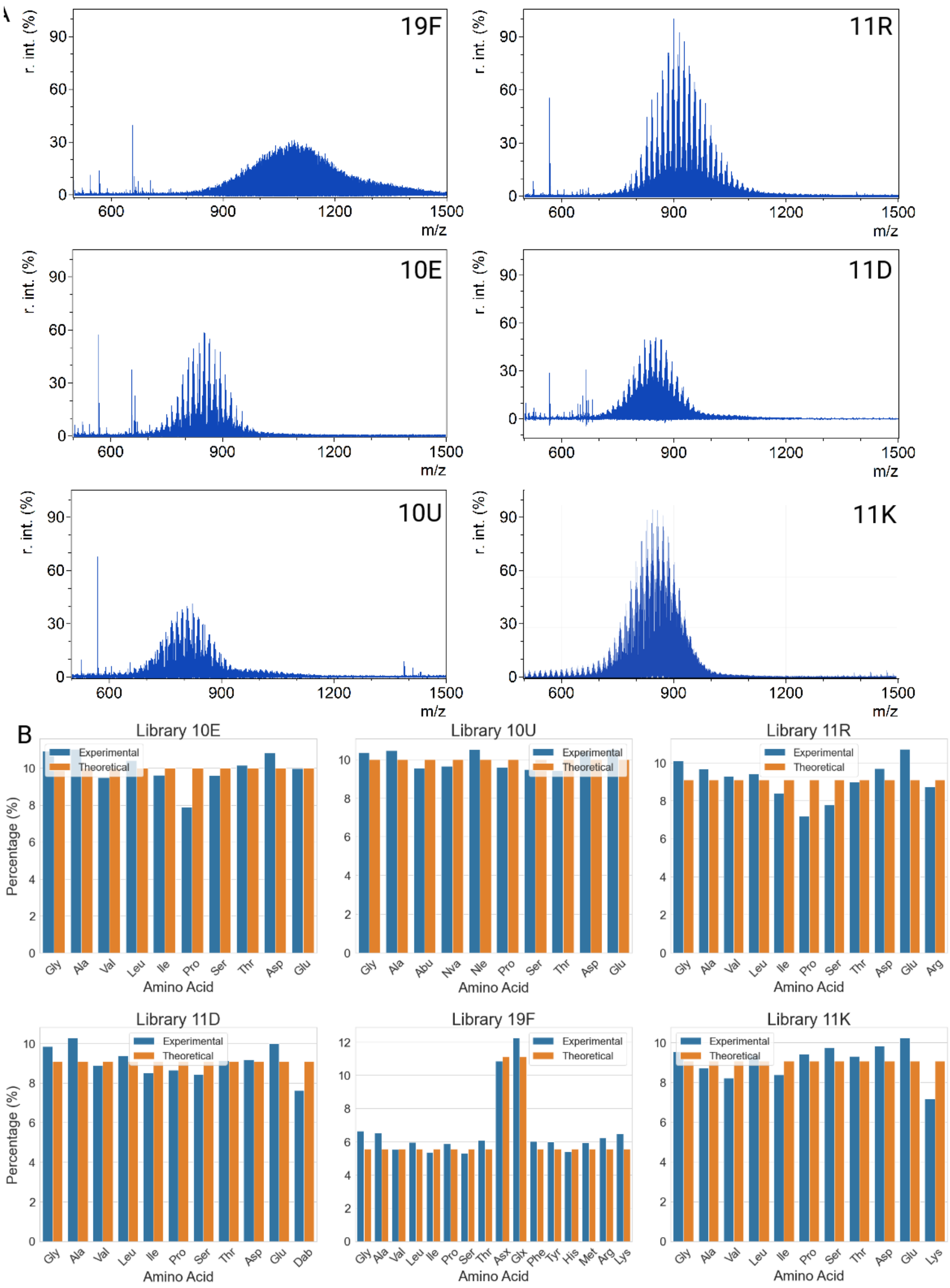


Figure S1. Matrix-assisted laser desorption/ionization (A) and HPLC amino acid analysis (B) of 8-mer random peptide libraries used in the current work, showing molecular size and amino acid distribution in a satisfactory range from theoretical ones. Since Asp and Glu overlap with Asn and Gln on the HPLC graph due to the method-associated hydrolysis of Asn to Asp and Gln to Glu, they are pulled together at the amino acid distribution graphs for 19F and labelled Asx and Glx, respectively. Data on Met was excluded from analysis due to the methodological limitations of the amino acid analysis.

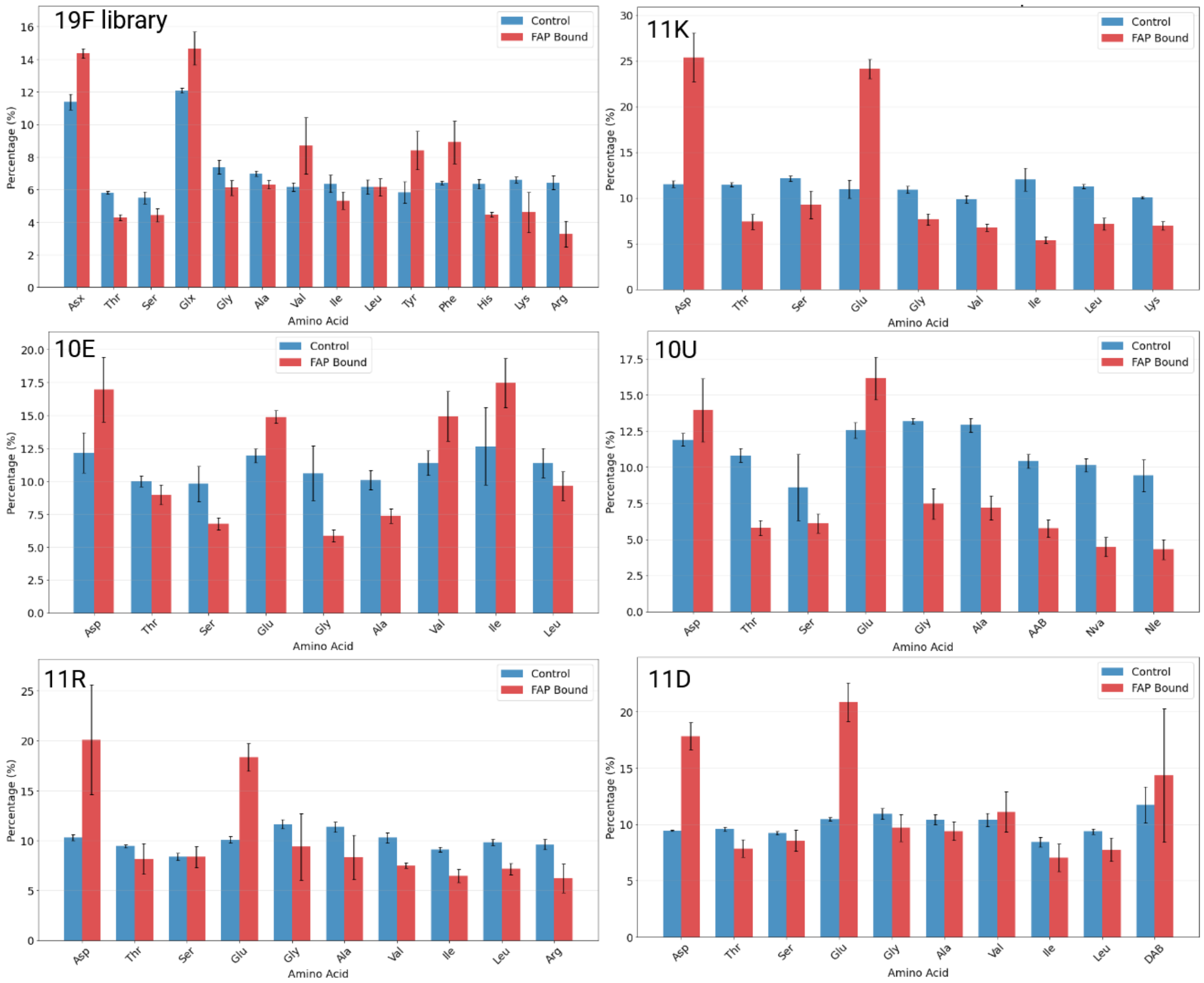


Figure S2. Amino acid distribution of random peptide libraries upon binding to fluorapatite. Each column represents a percentage share of individual amino acid in peptide composition (10% Gly = each $10^{th}$ residue in peptide library is glycine). We find enrichment in aspartate and glutamate in the bound fraction of every screened library, indicating their potential binding mediation. Since Asp and Glu overlap with Asn and Gln on the HPLC graph due to the method-associated hydrolysis of Asn to Asp and Gln to Glu, they are pulled together at the amino acid distribution graphs for 19F and labelled Asx and Glx, respectively.

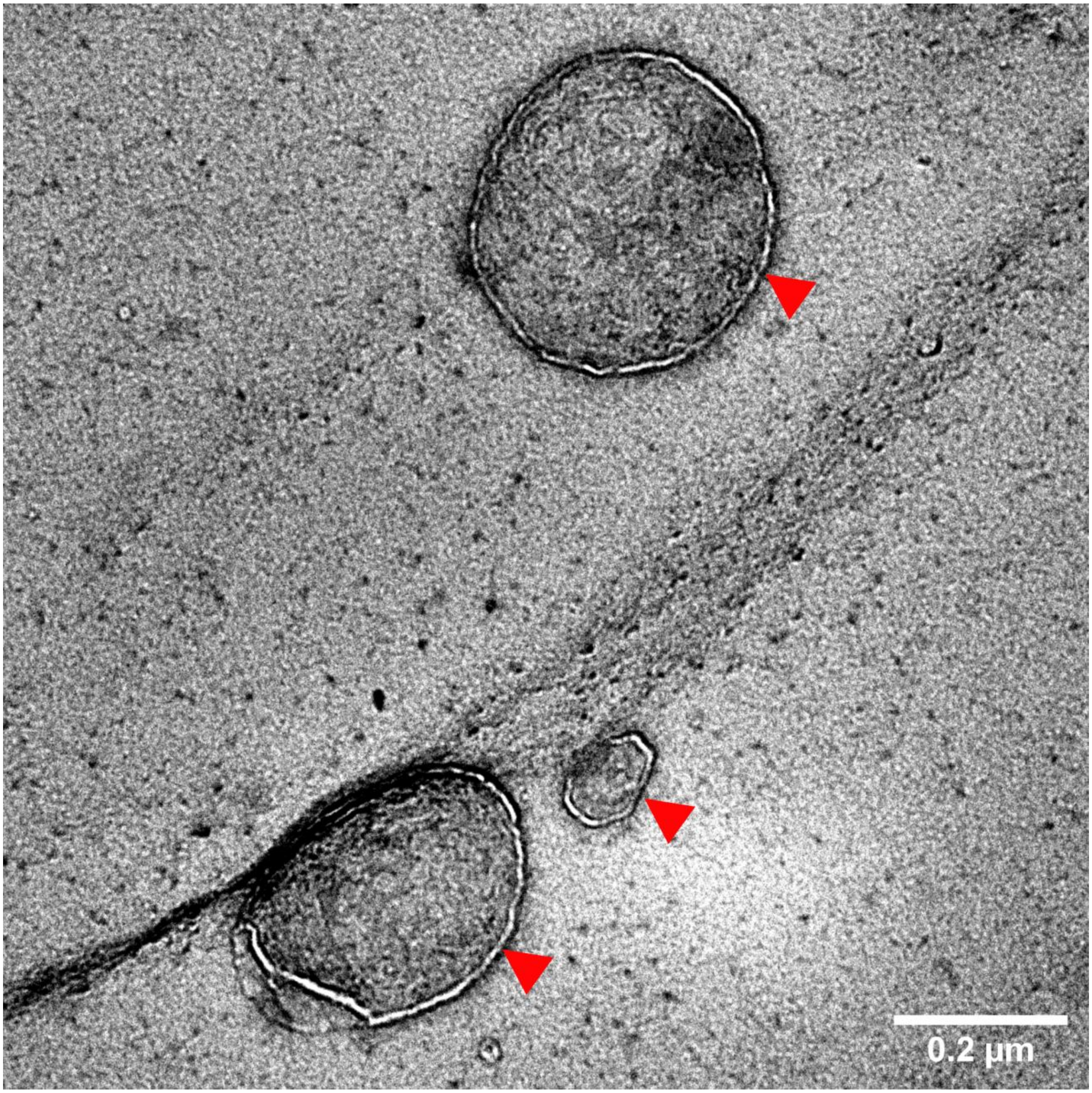


Figure S3. Transmission electron microscopy of decanoic acid:decanol (1:1) vesicles in PBS. Membranes are indicated by red arrows. Staining by uranyl acetate. Thickness of the membrane aligns with the expected range, and lumen is visible within a membrane. Scale bar - 200 nm.

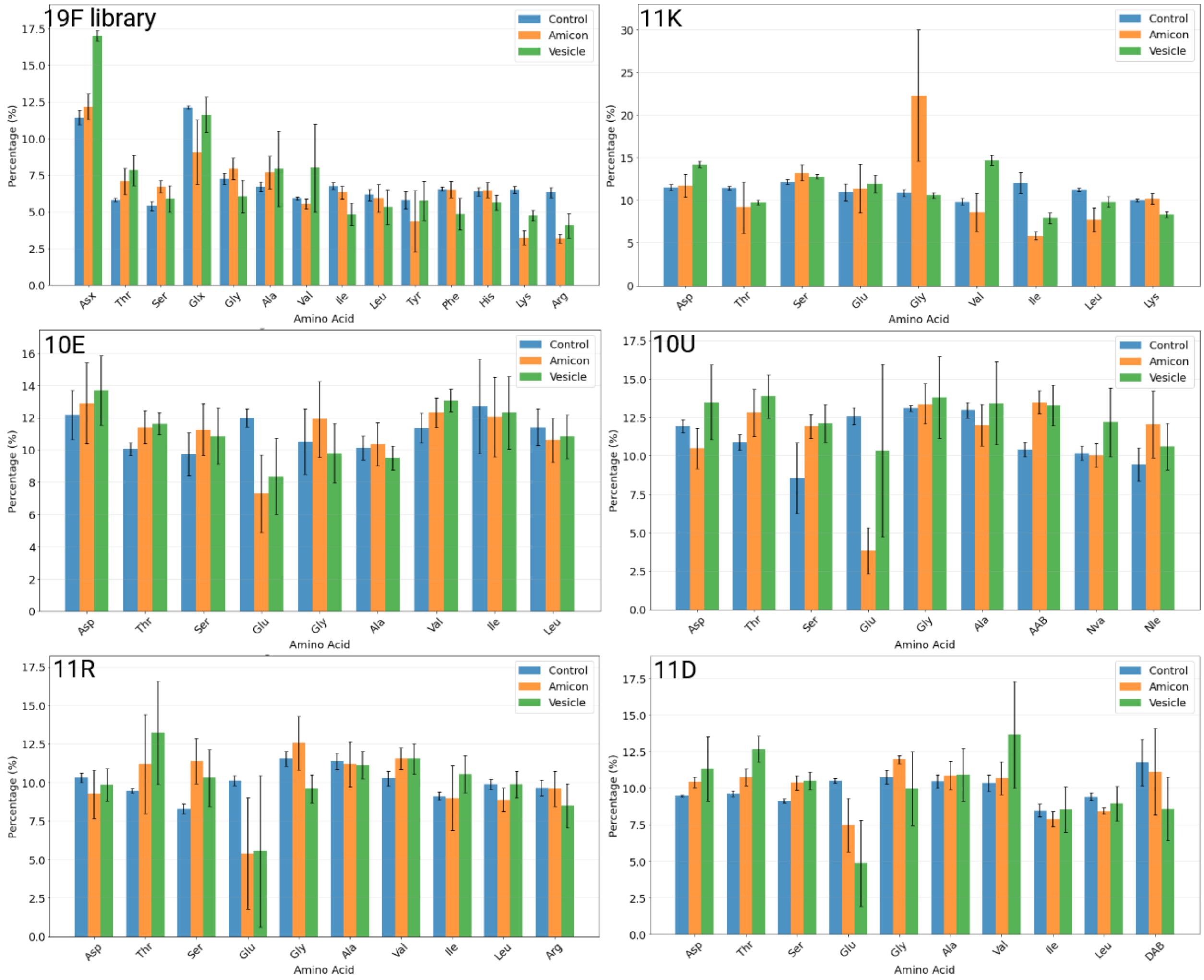


Figure S4. Amino acid distribution of random peptide libraries upon filtration through Amicon Ultra-4 3K centrifuge filters (orange bars) and upon incubation with DA:DOH vesicles with consequent filtration through Amicon Ultra-4 3K centrifuge filters (green bars). It appears that filtration through Amicon Ultra-4 3K filters mildly skews amino acid distribution of 8-mer peptide libraries, and, as such, it is difficult to derive conclusions on the changes of amino acid distribution of peptide library upon incubation with DA:DOH vesicles. Since Asp and Glu overlap with Asn and Gln on the HPLC graph due to the method-associated hydrolysis of Asn to Asp and Gln to Glu, they are pulled together at the amino acid distribution graphs for 19F and labelled Asx and Glx, respectively. Error bars represents standard deviation from six replicates.

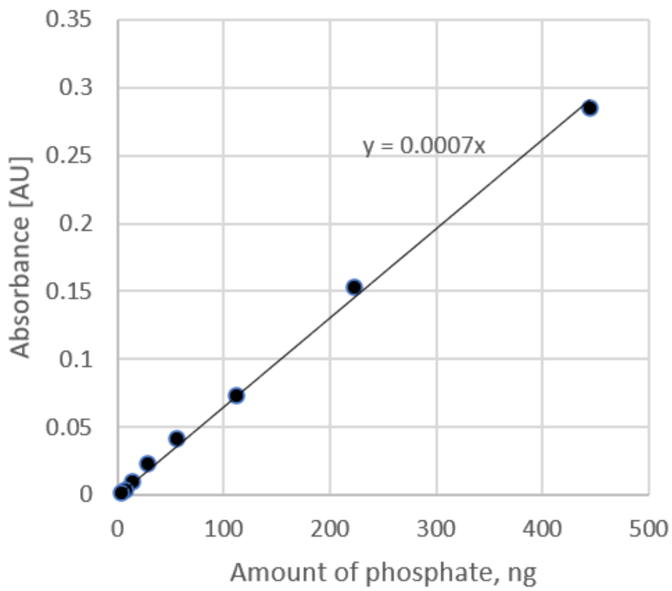


Figure S5. Calibration curve used for measurement of phosphate release from fluorapatite using molybdenum blue spectrophotometric assay. Absorbance values represent optical density at 882 nm. The working concentration range from 3.75 to 150 ng of phosphate (as $PO_4^{3-}$).

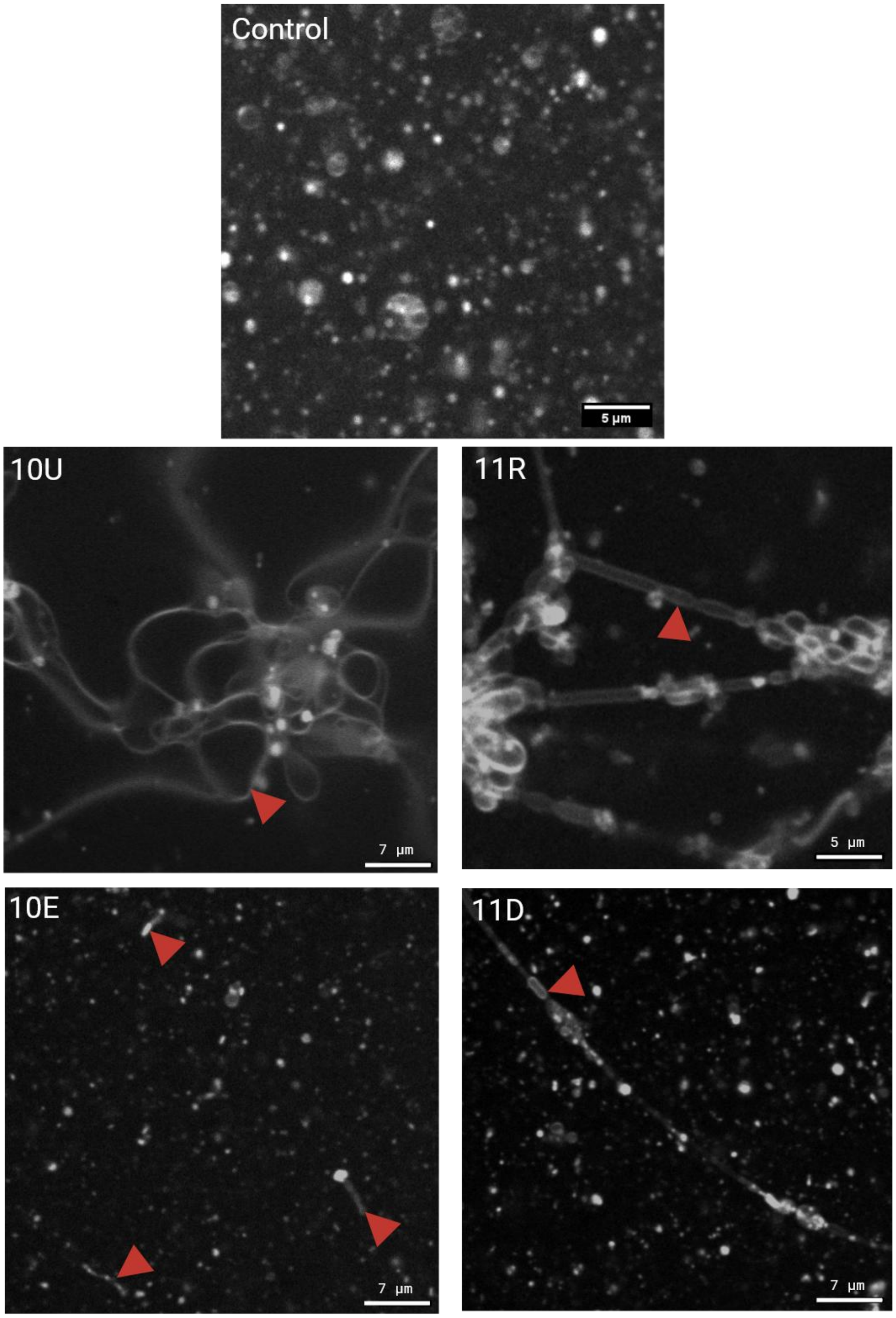


Figure S6. Detailed images of filamentous structures consistent with growing and budding membranes. With the exception of the 19F library, every peptide library screened induced membrane growth and budding upon addition of NaCl, but not in control conditions. Red arrows - growing membranes. Scale bar - 5 µm.

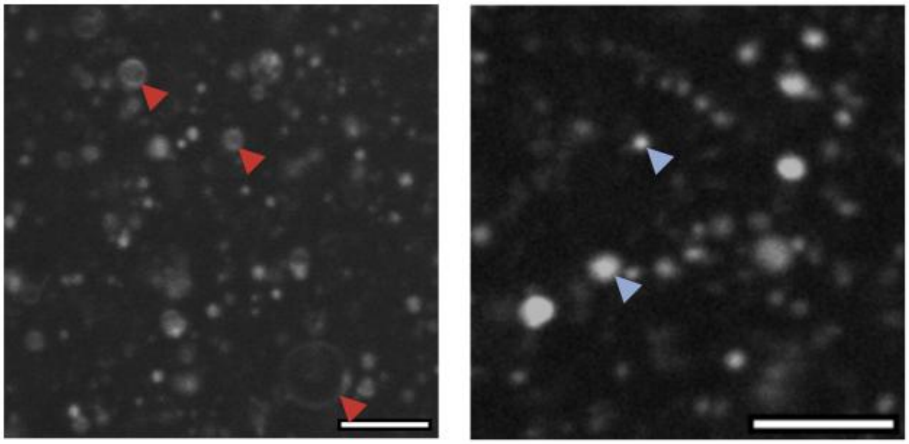

DA:DOH vesicles, pH 7.4, 60 s of vigorous shaking after addition of DOH, no shaking after addition of 1 % HCl

DA:DOH vesicles, pH 7.2, 60 s of vigorous shaking after addition of DOH + 60 s of vigorous shaking after addition of 1 % HCl

Figure S7. Example of methodological differences in vesicle stock preparation influencing vesicle stability. Observable differences in the sample composition as a result of relatively minor changes in handling suggests the need for greater caution when approaching the results quantitatively. Red arrow - unilamellar vesicles with visible lumen, blue arrow - punctate structures or oil droplets. Scale bar - 5 μm.

| | Control | Vesicles + 363 mM NaCl | Vesicles + 8.9 mM $MgCl_2$ | Vesicles + 10E library | Vesicles + 10E + NaCl | Vesicles + 10E + $MgCl_2$ |
|---|---|---|---|---|---|---|
| ζ-potential (mV) | -61.5±4.2 | -17.6±9.2 | -30.9±0.4 | -55.1±1.4 | -18.0±5.6 | -11.3±4.7 |
| Average particle size (nm) | 396.7±10.4 | N/A | N/A | 389.3±7.0 | N/A | N/A |
| Quality Factor | 2.0±0.5 | 0.8±0.6 | 1.3±.0.3 | 2.8±1.8 | 1.0±0.8 | 0.9±0.3 |

| | Vesicles + 19F library | Vesicles + 19F + NaCl | Vesicles + 19F + $MgCl_2$ | Vesicles + 10U library | Vesicles + 10U + NaCl | Vesicles + 10U + $MgCl_2$ |
|---|---|---|---|---|---|---|
| ζ-potential (mV) | -51.1±7.6 | -17.6±9.2 | -28.4±0.7 | -49.9±0.9 | -13.6±5.4 | -27.6±1.0 |
| Average particle size (nm) | 399.7±1.7 | N/A | N/A | 425.0±9.3 | N/A | N/A |
| Quality Factor | 2.0±0.3 | 0.8±0.6 | 1.3±0.4 | 1.3±0.2 | 0.3±0.1 | 1.2±0.1 |

| | Vesicles + 11R library | Vesicles + 11R + NaCl | Vesicles + 11R + $MgCl_2$ | Vesicles + 11D library | Vesicles + 11D + NaCl | Vesicles + 11D + $MgCl_2$ |
|---|---|---|---|---|---|---|
| ζ-potential (mV) | -52.1±0.8 | -5.0±2.5 | -20.3±3.3 | -52.7±0.5 | -15.9±1.7 | -26.5±0.6 |
| Average particle size (nm) | 415.3±9.5 | N/A | N/A | 395.9±17.2 | N/A | N/A |
| Quality Factor | 2.4±0.4 | 0.3±0.1 | 1.8±0.8 | 2.7±2.4 | 0.7±0.6 | 1.3±0.2 |

Table S1. Table presenting ζ-potential, particle size and conductivity of DA:DOH vesicles under different conditions. ζ-potential of DA:DOH vesicles drops upon addition of salt and selected peptide libraries, indicating their potential role in vesicle flocculation (in case of NaCl), collapse (in case of $MgCl_2$) and binding/functional effects (in case of peptide libraries). Particle size was not measured in certain conditions due to the particle size being out of the range of the instrument. Quality factor, as established by the Malvern Zetasizer instrument, represents signal to noise ratio of ζ-potential readings, with values above 0.9 considered to be of excellent quality and of 0.7-0.9 to be acceptable data with potential noise. We find higher noise upon addition of NaCl and $MgCl_2$, due to the increase in sample's conductivity and quick deterioration (corrosion) of the instruments' measuring cuvette by the electric current. Assay was not conducted on the 11K library.

**A. Phosphate release group comparison**

| Group 1 | Group 2 | p-value (Holm-Bonferroni corrected) |
|---|---|---|
| Control | 11D | 0.00848 |
| 11R | 11D | 0.008539 |
| 19F | 11D | 0.008539 |
| Control | 19F | 0.008599 |
| Control | 11R | 0. 008599 |
| Control | 10E | 0. 008599 |
| Control | 10U | 0. 008599 |
| 11R | 11K | 0.008658 |
| 19F | 11K | 0.008658 |
| 10U | 11K | 0.0022811 |
| 11R | 10E | 0.031048 |
| 10E | 11K | 0.031048 |
| 11D | 10U | 0.035529 |

**B. Fluorapatite binding group comparison**

| Group 1 | Group 2 | p-value (Holm-Bonferroni corrected) |
|---|---|---|
| 19F | 10E | 0.007194 |
| 19F | 11R | 0.033698 |

**C. Vesicle binding group comparison**

| Group 1 | Group 2 | p-value (Holm-Bonferroni corrected) |
|---|---|---|
| 19F | 10U | $8.77*10^{-7}$ |
| 11R | 10U | 0.00237 |
| 19F | 10E | 0.01337 |
| 11D | 10U | 0.03004 |

### D. Impact of peptide libraries and different salt conditions on $OD_{490}$ of a vesicle solution

#### 1. Control condition (no salts added to the solution)

| Group 1 | Group 2 | p-value (Holm-Bonferroni corrected) |
|---|---|---|
| 10E | 11R | 0.000003 |
| 10E | 19F | 0.000005 |
| 10E | 11D | 0.00018 |
| No peptide | 19F | 0.000186 |
| No peptide | 11D | 0.000256 |
| No peptide | 11R | 0.001303 |
| No peptide | 11K | 0.001851 |
| 10E | 11K | 0.002228 |
| No peptide | 10U | 0.037063 |
| 10E | 10U | 0.038740 |

#### 2. NaCl, 363 mM present

| Group 1 | Group 2 | p-value (Holm-Bonferroni corrected) |
|---|---|---|
| No peptide | 11D | 0.000001 |
| 10U | 11R | 0.000005 |
| No peptide | 11K | 0.000006 |
| No peptide | 10E | 0.000006 |
| No peptide | 10U | 0.000019 |
| No peptide | 19F | 0.000026 |
| No peptide | 11R | 0.000058 |
| 10U | 19F | 0.001758 |
| 10E | 10U | 0.003515 |
| 11D | 19F | 0.009529 |

**3. $MgCl_2$, 8.9 mM present**

| Group 1 | Group 2 | p-value (Holm-Bonferroni corrected) |
|---|---|---|
| No peptide | 11K | 0.009163 |
| No peptide | 10U | 0.012342 |
| No peptide | 11D | 0.012342 |
| 11R | 11K | 0.012342 |
| 11D | 11R | 0.021804 |

Table S2. Selected results of statistical analysis of values for phosphate release (A), fluorapatite binding (B), vesicle binding (C) and $OD_{490}$ (D) experiments. Only groups with a corrected p-value greater than 0.05 were included in the table.

| | Control (no libraries) | Vesicles + 19F library | Vesicles + 11R library | Vesicles + 10E library | Vesicles + 11D library | Vesicles + 11U library |
|---|---|---|---|---|---|---|
| DA+DOH concentration, mM | 384/373 nm ratio | | | | | |
| 0.354 | 1.08±0.02 | 1.12±0.002 | 1.09±0.003 | 1.10±0.004 | 1.09±0.003 | 1.09±0.008 |
| 0.708 | 1.13±0.03 | 1.11±0.004 | 1.08±0.01 | 1.10±0.004 | 1.08±0.005 | 1.11±0.004 |
| 1.416 | **1.19±0.01** | **1.18±0.02** | **1.19±0.03** | **1.19±0.03** | **1.15±0.02** | 1.14±0.02 |
| 2.832 | 1.26±0.01 | 1.21±0.01 | 1.24±0.03 | 1.27±0.02 | 1.23±0.02 | **1.22±0.008** |
| 5.664 | 1.32±0.01 | 1.28±0.02 | 1.34±0.02 | 1.35±0.01 | 1.33±0.02 | 1.30±0.004 |
| 11.327 | 1.40±0.019 | 1.37±0.01 | 1.42±0.001 | 1.42±0.01 | 1.42±0.01 | 1.38±0.007 |

Table S3. Changes in critical vesicle concentration (CVC) of DA:DOH in the presence of peptide libraries. Method used utilises changes in fluorescence of pyrene dye in the presence of vesicles. CVC was estimated to be at the 1.15 cutoff point of 384/373 nm ratio. In bold – the first 384/373 ratio that meets the CVC criterion. Data is average of triplicates with standard deviation. Assay was not conducted on the 11K library to eliminate redundancy.
